# Integrative Analysis of EGF, OSM, and TGFB Signaling Pathways Reveals Synergistic Mechanisms Driving Cell Motility Through CXCR2 Chemotactic Signaling and CREB Activation

**DOI:** 10.1101/2025.04.03.647095

**Authors:** Ian C. McLean, Sean Gross, Jeremy Copperman, Daniel Derrick, Laura Heiser

**Affiliations:** Department of Biomedical Engineering, OHSU, Portland, OR USA; Capsida Biotherapeutics Inc., Thousand Oaks, CA USA; Cancer Early Detection Advanced Research Center, OHSU, Portland, OR USA; Knight Cancer Institute, OHSU, Portland, OR USA

## Abstract

The microenvironment surrounding cells plays a critical role in determining cellular phenotype. Key components of the microenvironment include the diverse milieu of ligands and cytokines bind cell surface receptors to initiate changes in molecular programs. While the responses to extracellular signals have been extensively studied in isolation, little is known about the effects of combinations of signals on phenotypic and transcriptional responses. In this study, we used a coordinated approach to systematically investigate the combinatorial effects of the cytokines Oncostatin M (OSM) and Transforming Growth Factor Beta 1 (TGFB), and the growth factor Epidermal Growth Factor (EGF) on MCF10A mammary epithelial cells. Quantitative analysis of live-cell imaging data revealed a complex array of phenotypic responses after ligand treatment, including changes in proliferation, motility, cell clustering, and cytoplasmic size. We observed that all ligand combinations produce emergent phenotypic responses distinct from the maximal effects of individual ligands, indicating induction of new molecular programs. Companion RNA sequencing studies revealed a synergistic upregulation of a small but specific transcriptional program, including genes involved in cell migration, epithelial differentiation, and chemotactic signaling. Notably, these included chemokines such as CXCL3, CXCL5, and PPBP, which are known drivers of epithelial proliferation and migration. Additionally, transcription factor enrichment analyses and Reverse Phase Protein Array (RPPA) studies highlighted distinct changes in pathway utilization and transcription factor activity following combination treatment, including enhanced activation of MAP kinase and CREB signaling, compared to treatment with either agent alone. Using partial least squares regression, we identified robust transcriptional signatures associated with quantitative cellular phenotypes. We validated these signatures in independent datasets, confirming that they generalize across cellular contexts. Finally, an in-depth functional analysis of cell motility with RNA interference and pathway inhibition revealed that synergistic upregulation of CXCR2 signaling, mediated by CREB transcription factor activation, contributes to increases in cell motility across ligand conditions. These findings underscore the importance of combinatorial signaling in reprogramming epithelial phenotypes and reveal potential therapeutic targets for disrupting synergistic pathways in disease contexts such as cancer progression. Together, this study provides a framework for understanding how complex ligand interactions shape phenotypic and molecular landscapes.

## Introduction

The intricate interplay between cells and their microenvironment is a fundamental determinant of cellular behavior and function. Central to this dynamic relationship is the diverse array of ligands, cytokines, and extracellular matrix proteins that can initiate myriad intracellular responses that ultimately shape cellular phenotype. While the impacts of individual extracellular signals have been extensively studied, our understanding of how cells integrate and respond to combinations of extracellular signals remains limited.

In this study, we investigate the combinatorial effects of three ligands: Oncostatin M (OSM), Transforming Growth Factor Beta 1 (TGFB), and Epidermal Growth Factor (EGF), on the phenotypic and molecular responses of MCF10A mammary epithelial cells. These ligands hold pivotal roles in the normal development and function of mammary tissue, and their dysregulation is associated with disease (Hardy et al., 2010; Moses & Barcellos-Hoff, 2011; Tiffen et al., 2008). EGF canonically activates the MEK/ERK and PI3K signaling pathways (Herbst, 2004), OSM signals through JAK/STAT pathways (Dey et al., 2013), and TGFB orchestrates SMAD-mediated processes (Weiss & Attisano, 2013). The signaling pathways activated by EGF regulate the motility, proliferation, and invasion of normal and malignant breast epithelial cells (Herbst, 2004). TGFB induces epithelial-to-mesenchymal transition (EMT) in breast epithelial cells, achieved through the induction of EMT-associated transcription factors including SNAI1 and SNAI2, resulting in stereotyped changes in cell morphology and motility (Nawshad et al., 2005). TGFB also influences breast epithelial cell proliferation by activating p21 and suppressing key cell cycle transcription factors such as MYC, leading to cell cycle arrest (Donovan & Slingerland, 2000). Activation of JAK/STAT signaling downstream of OSM and other IL6 family cytokines modulates invasive properties and induces changes in modes of migration (Gross et al., 2022; Johnson et al., 2018). Despite their well-defined molecular consequences, the interplay between these signaling pathways and their combined impact on cellular behavior remains poorly understood.

Prior investigations into the combinatorial effects of perturbations have primarily focused on exploring and predicting the interplay of therapeutic inhibitors (Geary, 2013). Computational modeling approaches have demonstrated that synergistic drug interactions can be partially predicted from the transcriptional profiles of cells treated with individual agents (Bansal et al., 2014; Eslami et al., 2022; Lotfollahi et al., 2023; Niepel et al., 2017). In addition, there is evidence indicating a strong correlation between synergistic gene expression patterns and the degree of drug synergy, demonstrating a robust association between combinatorial transcriptional dynamics and resultant phenotypic responses (Diaz et al., 2020). However, the exploration of combinatorial effects and relationships between transcription and phenotype have typically been confined to a single phenotypic response, notably viability, without extending to more complex phenotypes such as cell motility, morphology or spatial arrangement. Expanding our understanding of combinatorial effects beyond drug perturbations and viability holds great promise for uncovering mechanisms of signal integration at molecular and phenotypic levels.

Employing live-cell imaging and RNA-seq, here we systematically investigate the phenotypic and transcriptional responses of MCF10A mammary epithelial cells to combinations of EGF, TGFB, and OSM. MCF10A is a well-characterized model system that has been utilized extensively to investigate tissue development, migration, and proliferation (Debnath et al., 2003; McQueen et al., 2018; Melani et al., 2008; Pierozan & Karlsson, 2018; Seton-Rogers et al., 2004). We found that all combinations of ligands induce phenotypic responses that differ significantly from their respective single ligand phenotypes, suggesting activation of additional molecular programs. Motivated by this finding, we performed comprehensive transcriptomic analysis to identify synergistic transcriptional programs in each combination condition, which revealed specific transcriptional programs modulated in response to combination treatments. We used Partial Least Squares Regression (PLSR) (Joreskog & Wold, 1982) to decipher the complex relationship between transcriptional programs and cellular phenotype. Our comprehensive analysis revealed that when combined, EGF and OSM synergistically amplify molecular programs associated with leukocyte chemotaxis and CXCR2 activation, resulting in increased cell motility. Functional validation demonstrated that synergistic upregulation of CXCR2-associated chemotactic factors is mediated by CREB transcription factor activation.

## RESULTS

### EGF, OSM and TGFB treatment combinations induce emergent phenotypic responses

We nominated for study a panel of three ligands (EGF, OSM and TGFB) that canonically activate distinct signaling pathways and that have been shown to induce strong phenotypic responses in MCF10A mammary epithelial cells (Gross et al., 2022) (Supp Figure 1A) **(Figure 1A)**. Although the effects of these individual ligands have been examined in various cellular contexts including MCF10A (Gross et al., 2022; Kim et al., 2004, 2005; Sundqvist et al., 2020; Tarcic et al., 2012), predicting how cells will respond to dual treatment at the molecular or phenotypic level is challenging due to the complex responses elicited by single ligands.

**Figure 1:**
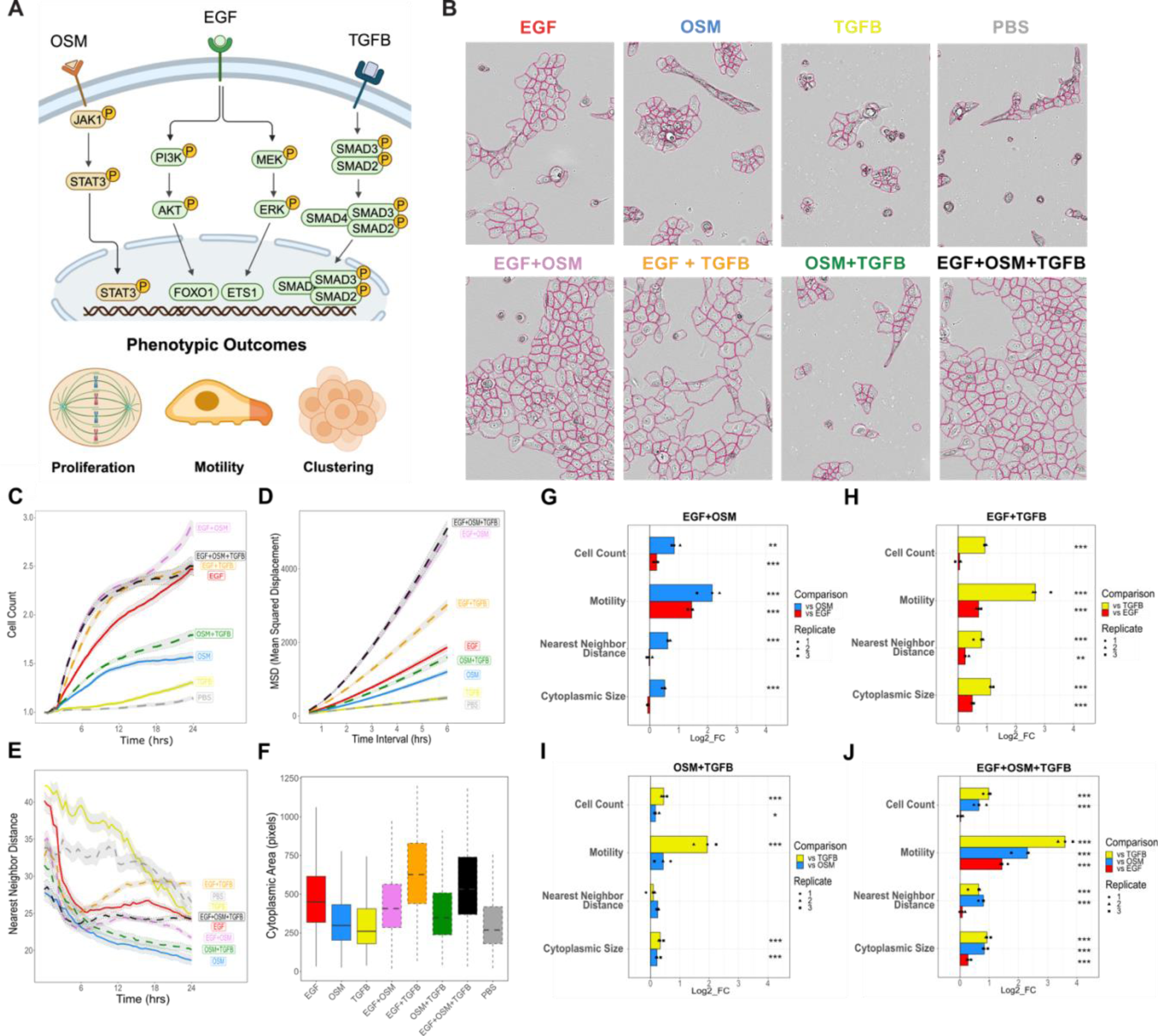
Combination treatments induce diverse and emergent phenotypic behavior. MCF10A cells were treated with PBS (control), EGF, OSM, or TGFB alone, or their pairwise/triple combinations, and subjected to live-cell imaging and quantitative analysis over 24 hours. (A) Overview of the signaling pathways activated by each treatment, highlighting the canonical pathways associated with each ligand. Schematic representations depict the potential phenotypic outcomes (e.g., proliferation, motility, and clustering/spreading) influenced by these pathways. (B) Representative images of MCF10A cells under different ligand treatments at 24 hours, demonstrating changes in cell phenotype. (C-F) Quantification of cell phenotype from 0-24 hours (cell count normalized to T0, MSD, nearest neighbor distance) or at 24 hours (cytoplasmic area). (G-J) Quantified phenotypic responses for each combination condition were compared to each single ligand condition comprising that combination. ANOVA followed by post-hoc Tukey’s honest significant difference test was used to assess significance, with p-value < 0.05 considered significant.

MCF10A cells cultured in growth factor free media were exposed to EGF, OSM, and TGFB independently, or in pairwise or triple combination, and subjected to live-cell imaging every 30 minutes for 24 hours. EGF is typically included as a supplement in MCF10A culture media, so this condition serves as a positive control for our study (Herbert D. Soule, 1990); PBS serves as a growth-factor-starved negative control. We employed quantitative image analysis to assess changes in cell count, motility, cell clustering and cell spreading, and cytoplasmic size (**Figure 1B**). In our initial analyses, we describe the quantitative responses to each single ligand compared to PBS control. Consistent with previous studies (Gross et al., 2022), we found that independent treatment with EGF strongly induced proliferation, whereas TGFB and OSM induced only modest increases in cell count compared to PBS **(Figure 1C, Supp Fig 1A)**. Cell motility was evaluated by calculating the mean squared displacement (MSD) over all cells for each treatment and deriving the diffusion coefficient from the MSD curves **(Figure 1D)**. Similar to cell proliferation, EGF treatment significantly increased cell motility compared to PBS control, while OSM induced a modest increase in motility, and TGFB showed no change in motility compared to PBS **(Figure 1D, Supp 1A)**. The calculation of the distance to the nearest neighboring cell revealed changes in cell clustering (decreased neighbor distance) and cell spreading (increased neighbor distance). EGF and TGFB treatment did not significantly affect the nearest neighbor distance compared to PBS control; however, OSM led to a decrease in this metric, indicating tight cell clustering **(Figure 1E, Supp Fig 1A)**. Lastly, we observed that EGF treatment led to an increase in cytoplasmic size compared to the PBS control, whereas TGFB or OSM alone had no effect **(Figure 1F, Supp Fig 1A)**. Notably, the addition of TGFB to EGF further amplified the increase in cytoplasmic size beyond that induced by EGF alone **(Supp Fig 1B)**. In total, these analyses demonstrate that each treatment regime induced a distinct constellation of phenotypic changes.

Given the distinct phenotypic responses associated with each ligand, we hypothesized that the phenotypic responses to combination treatments would adhere to the Highest Single Agent (HSA) model (Berenbaum, 1989). The HSA model, used to study drug combination synergy, posits that the expected phenotypic response for a combination treatment is equal to the maximal effect of any single agent. Using this framework, we anticipated that the phenotypic responses to combination ligand conditions would primarily reflect the influence of the single ligand with the most significant impact on each aspect of cell phenotype. To test this hypothesis, we compared the quantified phenotypic metrics for each combination condition to those of each respective single ligand condition. A phenotype was considered emergent (i.e., it did not mirror either single ligand response) if we detected a significant change that deviated from each single ligand condition, assessed using a one-way analysis of variance (ANOVA) followed by post-hoc Tukey’s honest significant difference test (p-value < 0.05).

Consistent with the HSA model, cell count observed under EGF+TGFB and EGF+OSM+TGFB treatments remained unchanged compared to EGF alone. However, EGF+OSM and OSM+TGFB treatments resulted in a cell count greater than their individual ligand treatments **(Figure 1G, H, I, J).** This finding suggests an interaction between OSM and the other ligands that influences changes in cell proliferation. We also observed emergent phenotypic changes in cell motility, distance to nearest neighboring cells, and cytoplasmic size when ligands were applied in combination. Treatment with OSM and TGFB individually resulted in decreased cell motility when compared to EGF treated cells (**Supp Fig 1B**). However, combination treatment with EGF+OSM, EGF+TGFB, and EGF+OSM+TGFB resulted in an emergent increase in cell motility compared to all single ligand conditions (**Figure 1G, H, J**). Finally, combination treatment with EGF+TGFB resulted in an emergent phenotypic increase in both nearest neighbor distance and cytoplasmic size, indicating cell spreading (**Figure 1H**). Our findings indicate that across various ligand combination treatments and multiple phenotypic responses, combination treatment induces emergent behavior that deviates from that of individual ligand treatments. This suggests that combination treatments induce molecular programs not induced by single ligand treatments.

### Identification of molecular programs induced by single and combination ligand treatments

We next used RNA sequencing to examine the molecular mechanisms driving response to ligand combinations. Cells were harvested after 24 hours of ligand treatment and subjected to bulk RNA sequencing. We posited that treatments leading to the strongest changes in cell phenotype compared to the PBS control would likewise display the greatest change in transcriptional responses. To test this hypothesis, we first assessed the overall transcriptional perturbation for each ligand condition by quantifying the number of differentially expressed genes as compared to PBS control (LFC > 1.5 or LFC < -1.5 and q-value < .05).

Comparison of differentially expressed genes revealed that treatments modulate gene expression programs to different extents; the EGF+OSM treatment induced the greatest number of differentially expressed genes, while independent treatment with TGFB had only modest impact on transcription (**Figure 2A, Supp Fig 2A, 2B**). To assess the overall change in cell phenotype for each condition, we calculated the total magnitude of the four phenotypic metrics relative to the PBS control. This was done by representing the phenotypic metrics as a four-dimensional vector, where the origin indicates no change from PBS control, and then computing overall change in cell phenotype as the vector magnitude across these four dimensions. Then, for each treatment, we directly compared the overall change in cell phenotype to the number of differentially expressed genes (**Figure 2B**). We found a strong correlation between number of differentially expressed genes and overall change in phenotype (R^2^ = 0.75, p-value = .012). The treatments that provoked the most significant changes in gene expression also corresponded to the most substantial overall changes in cell phenotype (EGF+OSM+TGFB, EGF+OSM, and EGF+TGFB). Conversely, treatments that exerted a minimal change on cell phenotype led to the fewest changes in gene expression, highlighting a clear relationship between gene expression alterations and phenotypic outcomes across different treatment conditions.

**Figure 2:**
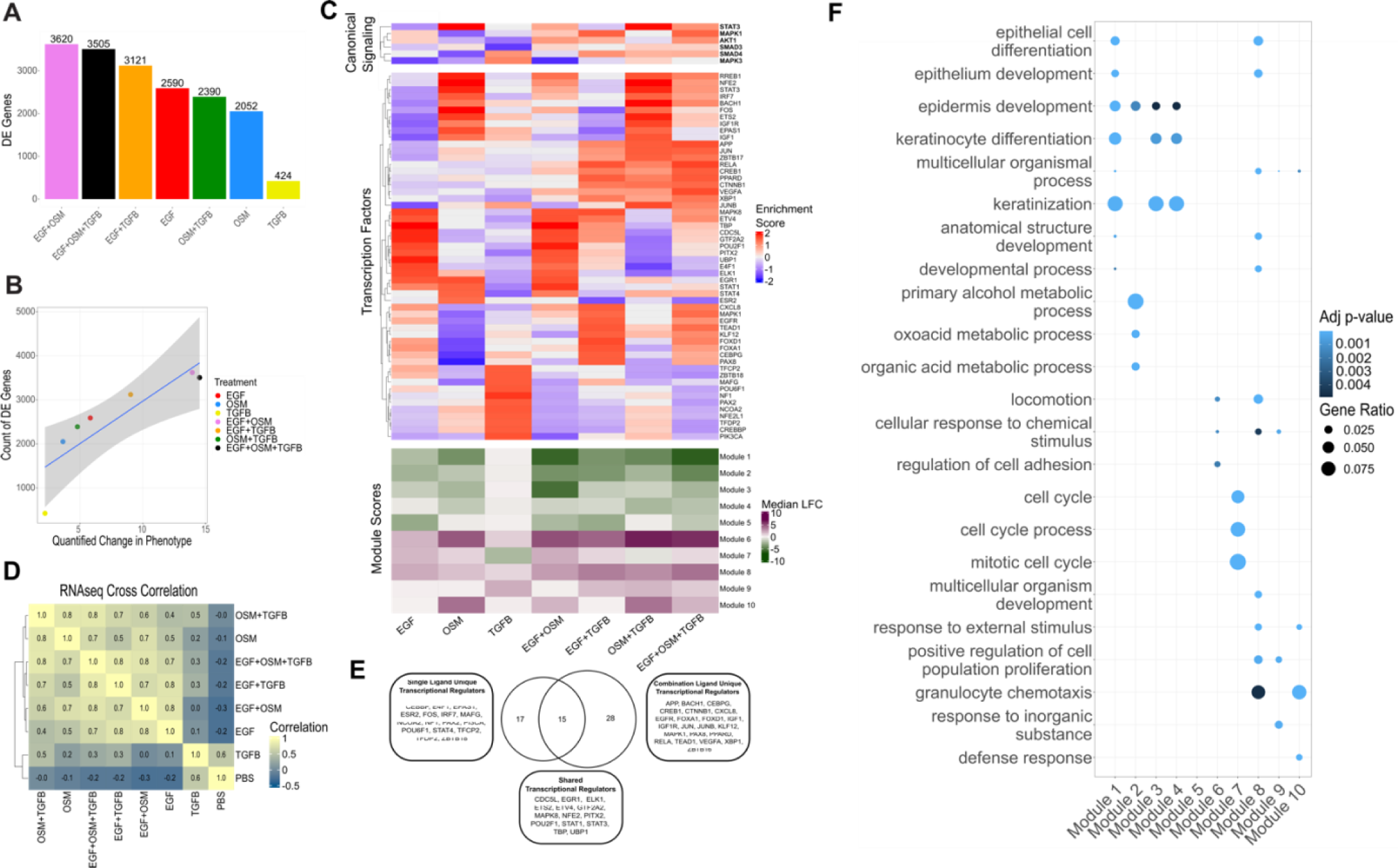
Transcriptional programs induced by single and combination ligand treatments. (A) Number of differentially expressed genes (LFC > 1.5 or < -1.5, q-value < 0.05) across treatments relative to PBS control, showing the greatest response in EGF+OSM treatment. (B) Correlation between the number of differentially expressed genes and overall changes in cell phenotype (R² = 0.75, p = 0.012). (C) Enrichment analysis of canonical transcriptional regulators (Canonical Signaling, top heatmap), the top decile of transcription factors (Transcription Factors, middle heatmap), and median LFC for gene modules (Module Score, bottom heatmap) activated in response to each treatment condition. (D) Pairwise Pearson correlations of gene expression log fold changes compared to T0 between all treatments. (E) Venn diagram comparing transcriptional regulators enriched in single versus combination treatments. (F) Enrichment of Gene Ontology terms for each gene module identified.

To further investigate the transcriptional programs modulated by single and combination treatments, we next performed transcription factor enrichment on the RNAseq dataset using Priori, an approach that leverages prior biological information to infer the activity of 223 transcriptional regulators (Yashar et al., 2024). Priori evaluates the activation levels of canonical transcription factors and enables data-driven identification of key signaling proteins that may play a crucial role in mediating phenotypic and combination responses. Analysis of the relative enrichment for canonical transcriptional regulators associated with EGF, OSM, and TGFB signaling with single-ligand treatments recapitulates established signaling pathways. Namely, EGF enriches AKT and MAPK transcriptional processes, OSM induces STAT3, while TGFB induces SMAD4 programs, respectively (**Figure 2C, Canonical Signaling**). Relative to EGF alone, EGF+TGFB increases MAPK enrichment, EGF+OSM increases AKT enrichment, and the three-ligand combination condition increases enrichment for both pathways. This suggests that activation of the MAPK and PI3K signaling cascade may contribute to the high level of proliferation observed in these conditions. Conversely, EGF+OSM diminishes STAT3 enrichment relative to the OSM condition (**Figure 2C, Canonical Signaling**). We also compared the top decile of transcription factors most strongly induced by each treatment condition **(Figure 2C, Transcription Factors)**, which revealed that approximately half of the transcriptional regulators enriched in single ligand treatments are also active in their respective combination conditions (16/33) (**Figure 2E, Supp Fig 2C**). However, 28 (28/44) transcriptional regulators show enrichment only in combination treatments and not in single ligand conditions. These factors comprise a diverse array of transcriptional regulators including growth factor signaling molecules (EGFR, IGF1, IGF1R, MAPK1, VEGFA) and regulators of epithelial and immune cell differentiation (APP, BACH1, CEBPG, CREB1, CTNNB1, RELA). This may indicate that a common molecular program is induced in response to treatment with multiple ligands.

We next sought to identify coordinated molecular programs associated with the observed phenotypic responses. Focusing on the subset of genes most strongly induced relative to the T0 control, we filtered for the top 200 most up-regulated and top 200 most down-regulated genes for each treatment (p-value < 0.05, LFC > 1.5). Pairwise correlation of the log fold changes revealed that the combination treatments share transcriptional similarities with at least one of the individual ligands in each pair (Pearson correlation > 0.7) (**Figure 2D**). To further explore what may be driving the distinct phenotypic responses in the combination conditions, we applied K-means clustering (Lloyd, 1982) and gap analysis (Tibshirani et al., 2001), which identified 10 coordinated gene modules (**Supp Fig 2D**). Gene module scores, calculated as the median expression for each module, were not exclusively up-or downregulated in any combination condition suggesting that combination responses do not activate unique transcriptional gene modules distinct from single ligand conditions, but rather modulate existing transcriptional programs (**Figure 2C, Module Scores**).

We further explored the molecular programs in each module through gene set enrichment analysis, focusing on curated Gene Ontology categories (**Figure 2F**) (Aleksander et al., 2023; Ashburner et al., 2000; Kolberg et al., 2023). Gene modules 1, 2, 3, and 4 are enriched for terms related to epithelial cell and keratinocyte differentiation and development. These modules are downregulated across all treatments except for the TGFB condition. Additionally, Modules 1 and 2 are more strongly downregulated by the EGF+OSM and EGF+OSM+TGFB conditions as compared to their respective single ligand treatments (**Supp Fig 2E, 2H**). These two treatments also show the most significant increase in motility compared to the PBS control, suggesting that downregulation of these gene modules corresponds to dedifferentiation associated with cell motility (**Figure 1C, 1D**).

Module 7 is enriched for cell cycle-related terms, with its expression highest under the EGF and EGF+OSM conditions, consistent with the increased cell count observed in these conditions. Gene Ontology terms associated with general cell motility (locomotion) are significantly enriched in modules 6 and 8, while specific mechanisms of cell motility (granulocyte chemotaxis) are enriched in modules 8 and 10. Module 6 is upregulated by EGF+TGFB, OSM+TGFB, and EGF+OSM+TGFB compared to all single ligand responses, suggesting an underlying amplification of general cell motility transcriptional programs when these signals are combined **(Supp Fig 2F, 2G, 2H)**. Conversely, module 10 is only upregulated by conditions containing OSM, indicating that a unique chemotaxis program is activated only when OSM is included in the treatment.

### Combination treatments result in specific synergistic transcriptional programs

The previous analyses suggest that combination treatments primarily amplify the transcriptional programs induced by single ligand responses. To further investigate this observation, we compared the transcriptional responses in combination treatments to predictions made using a simple additive model. Previous studies have shown that upregulated gene responses to ligand combinations typically follow either additive or multiplicative patterns (Sanford et al., 2020). To determine if this applies to our dataset, we developed a simple model by summing the fold changes in expression relative to the T0 control for upregulated genes (LFC > 0.5) from single ligand conditions and compared these predicted values to the observed expression levels in the combination treatments. Using this model, our analysis revealed a strong correlation between the predicted and actual responses for all ligand combination conditions (EGF+OSM - R² = 0.73, p-value < .001; EGF+TGFB - R² = 0.70, p-value < .001; OSM+TGFB - R² = 0.76, p-value < .001; EGF+OSM+TGFB - R² = 0.66, p-value < .001), suggesting that the combination responses can largely be explained by a simple additive model **(Supp Fig 3A-D)**. These findings indicate that ligand combination responses predominantly recapitulate the transcriptional responses of the individual ligands.

The observation that the transcriptional responses to combination treatments predominantly reflect the effects of individual ligands was unexpected, considering the complex phenotypic changes we observed in each combination condition. We hypothesized that these emergent phenotypic effects might arise from specific synergistic transcriptional patterns not captured in our previous analysis. Here we more deeply explore that observation by borrowing frameworks developed in drug combination studies to identify synergistic gene expression programs by applying HSA modeling to molecular quantification (Diaz et al., 2020). For each combination treatment, we calculated the log fold change in expression for all genes compared to each respective single ligand condition, then visualized the changes in expression in x-y scatterplots (**Figure 3A-C**). We designated a gene as positively synergistic if the log fold change in expression after combination treatment exceeded 1.5 compared to each of the single ligand conditions (adjusted p-value < 0.05) (**Figure 3A-C**). Similarly, we defined negatively synergistic genes as those with a log fold change of at least -1.5 compared to both respective single ligand conditions. This comprehensive analysis revealed that for each pairwise combination, there exist subtle yet significant synergistic transcriptional programs. The number of positively synergistic genes per combination condition ranged from 17 to 110, while negatively synergistic genes ranged from 27 to 148. To evaluate how the range of synergistic modulation in our ligand combinations compares to previous studies using drug combinations, we performed the same analysis on MCF7 malignant breast epithelial cells treated with Tamoxifen, Mefloquine, and Withaferin individually and in combination **(Supp Fig 3E-G)** (Diaz et al., 2020). These drugs were selected based on their phenotypic synergy in reducing cell viability when combined. When comparing our results, we found that although the degree of transcriptional synergy in our ligand combinations was far surpassed by the level of synergy observed in the Mefloquine and Tamoxifen combination (2235 genes), it was comparable to the two other combination conditions (50 genes, 137 genes). Interestingly, even in the Mefloquine and Withaferin combination, which showed relatively small transcriptional synergy, there was still substantial phenotypic synergy. This suggests that while a limited number of genes may be synergistically expressed in our ligand combination treatments, their strong modulation may have functional importance.

**Figure 3:**
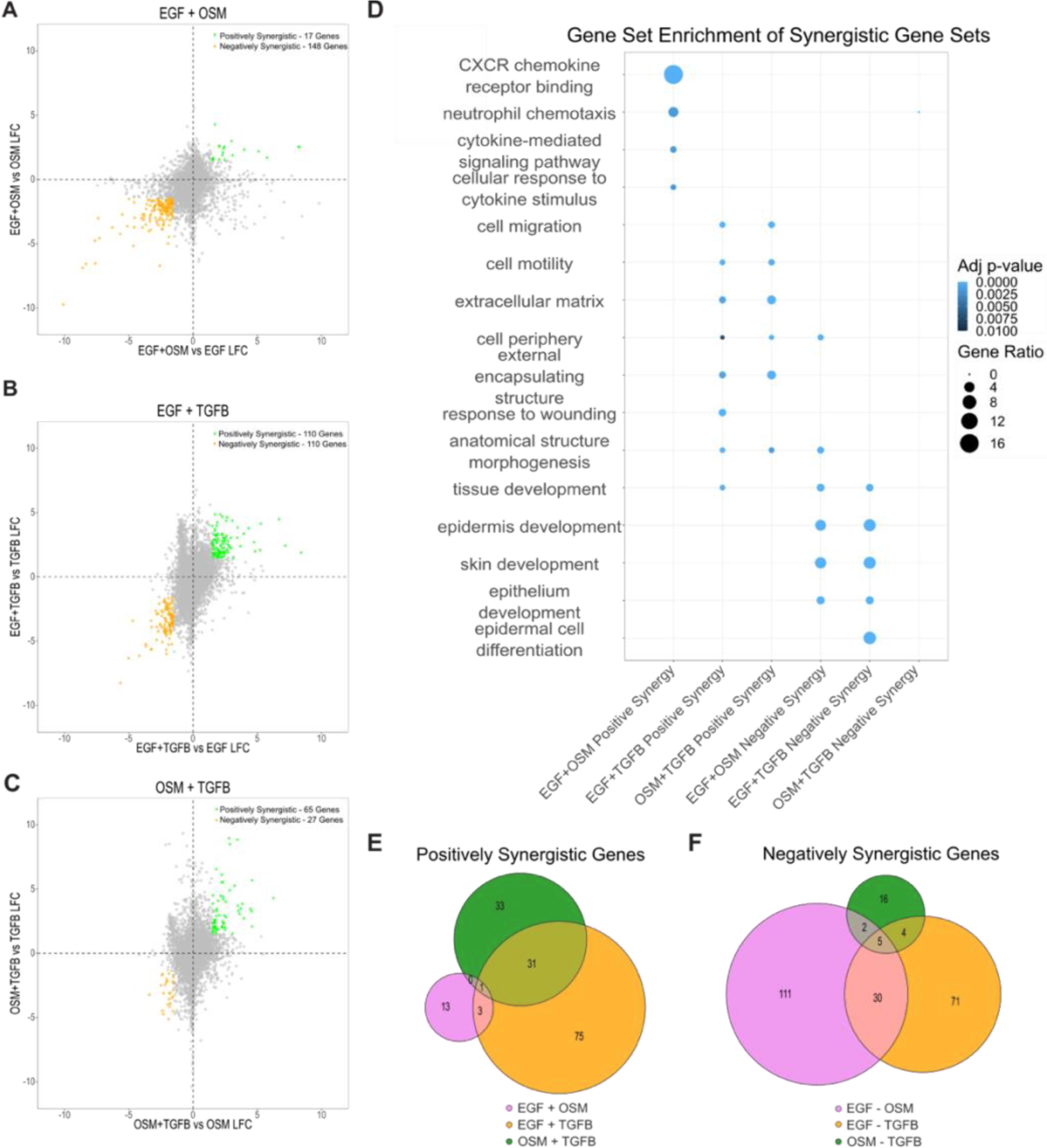
Synergistic transcriptional programs induced by ligand combination treatments. (A-C) Scatterplots showing log fold change in gene expression for combination treatments versus each respective single ligand condition (EGF+OSM, EGF+TGFB, OSM+TGFB). Genes exhibiting positive synergy (LFC > 1.5, adjusted p-value < 0.05 compared to both treatments) are shown in green, and genes with negative synergy (LFC < -1.5, adjusted p-value < 0.05 compared to both treatments) are shown in orange. (D) Gene Ontology enrichment analysis of positively and negatively synergistic gene sets. (E) Venn diagram showing overlap of positively synergistic genes between combination conditions. (F) Venn diagram depicting negatively synergistic genes between combination conditions.

We performed gene set enrichment analysis of the positive and negative synergistic gene sets identified for each combination treatment using Gene Ontology Biological Process, Molecular Function, and Cellular Component gene sets (Aleksander et al., 2023; Ashburner et al., 2000; Kolberg et al., 2023). Positively synergistic genes induced by the combination of EGF and OSM resulted in the upregulation of chemotactic transcriptional programs (**Figure 3D**). Combinations containing TGFB exhibited a large overlap of positively synergistic genes (EGF+TGFB = 32/110, OSM+TGFB = 32/65) **(Figure 3E)**. These combinations also synergistically induced transcriptional programs related to ECM remodeling and cell motility, consistent with TGFB’s known role in promoting epithelial-to-mesenchymal transition and the emergent enlargement of cell cytoplasm in this condition (Xu et al., 2009). Notably, these EMT-related programs were not strongly induced by TGFB treatment alone and required either OSM or EGF in combination. This aligns with the observation from live-cell imaging that TGFB only induces migration and alterations in cell morphology in the presence of OSM or EGF **(Figure 1D, E**). Consistent with the live-cell image data, EGF+OSM and EGF+TGFB both downregulate epithelial differentiation processes (epithelium development, skin development). Epithelial cells that undergo transdifferentiation and loss of epithelial identity, notably through epithelial-to-mesenchymal transition, exhibit increased motility and changes in cell morphology, which we observed in both treatment combinations (Dong et al., 2009). Negatively synergistic gene sets for EGF+OSM and EGF+TGFB had a large overlap, with 35 genes in common **(Figure 3F)**, suggesting that these shared repressive transcriptional programs contribute to these processes **(Figure 1D, E)**.

### Partial Least Squares Regression uncovers transcriptional signatures driving cellular phenotype

We next sought to more directly link image and RNAseq data to uncover molecular drivers of complex cellular phenotypes. To achieve this goal, we utilized Partial Least Squares Regression (PLSR), an efficient statistical prediction tool that is especially appropriate for small sample data with many possibly correlated variables (Joreskog & Wold, 1982). We constructed PLSR models to identify gene signatures to predict each of the four phenotypic metrics derived from live-cell imaging data (**Figure 1**). The three biological replicates of imaged cells shown in Figure 1 were harvested for the RNA sequencing described above, enabling direct linkage of image and RNAseq data for each experimental sample. This design leveraged biological variation in both phenotypic and transcriptional responses across replicates. The inputs for each model consisted of replicate-level phenotypic metrics and replicate-level Log Fold Change (LFC) gene expression compared to T0 control. Leave-one-out analysis was used to determine the optimal number of latent variables in each model and to evaluate model robustness (Hastie et al., 2009). The Relative Root Mean Squared Error of Prediction (RRMSEP) for the four models ranged from 0.205 to 0.563, indicating good fit.

For each PLSR model, we identified the gene signature most strongly associated with phenotypic changes by calculating Variable Importance in Projection (VIP) scores (Wold et al., 2001). VIP scoring estimates the overall significance of each feature in the model without distinguishing between positively and negatively correlated features. To further refine the top-scoring genes in each model, we categorized them based on their correlation with the phenotype of interest. For each model, we selected for further analysis the top 100 genes with the highest VIP scores and a positive correlation to the model’s first component and the top 100 genes with a negative correlation to the first component. There were varying degrees of overlap among the highest-scoring VIPs from different phenotypic signatures. The largest overlap was observed between the Nearest Neighbor Distance and Cytoplasmic Size signatures, likely due to the inverse relationship between cell spreading and cell clustering (**Fig 4A, B**). Similarly, there was overlap between the Cell Count and Motility signatures, which is expected as these biological processes have been shown to be driven by similar molecular mechanisms (Nair et al., 2019). Despite these overlaps, a significant number of genes were uniquely identified as VIPs for each phenotype, indicating that the PLSR regression identifies distinct biological mechanisms associated with each phenotype. We conducted gene set enrichment analysis for each signature to investigate the underlying cellular processes (Aleksander et al., 2023; Ashburner et al., 2000; Kolberg et al., 2023). Enrichment for the top negatively correlated VIPs for Nearest Neighbor Distance include terms related to inflammatory response (acute-phase response, complement activation) (**Supp Fig 4A, B**). The top positively correlated VIPs associated with cytoplasmic size are enriched in terms related to synthesis processes for cell membrane components (isoprenoid biosynthetic process, cholesterol biosynthetic process, phospholipid biosynthetic process) and cell-substrate interactions (cell-substrate junction, focal adhesion), indicating a shared transcriptional program involved in membrane and cytoskeletal remodeling that regulates cytoplasmic size **(Supp Fig 4C, D)**.

**Figure 4:**
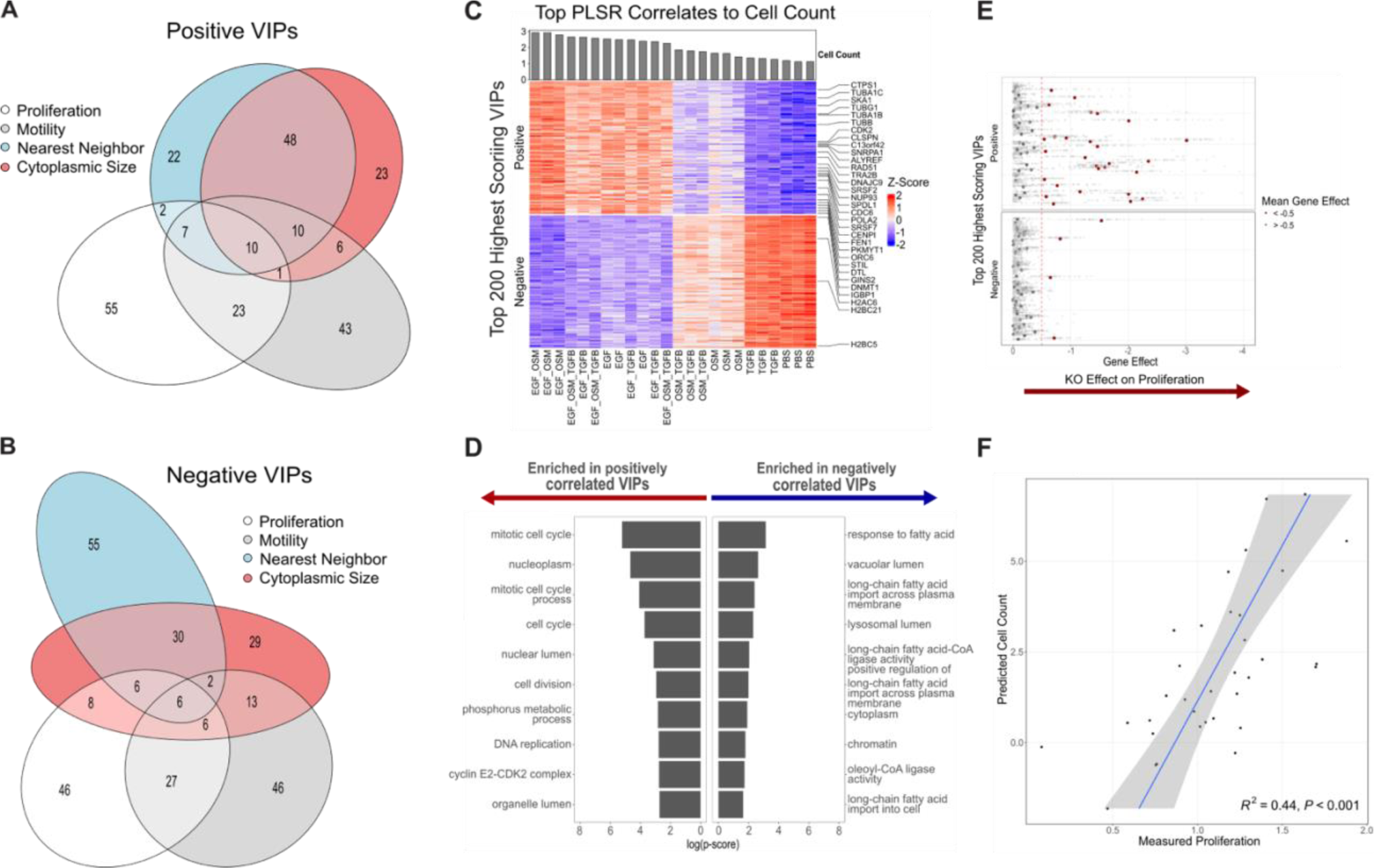
Partial Least Squares Regression (PLSR) links transcriptional signatures to phenotypic metrics. (A-B) PLSR models were used to link RNA sequencing data to phenotypic metrics derived from live-cell imaging (Cytoplasmic Size, Nearest Neighbor Distance, Motility, and Cell Count). Overlap in top positive VIP genes for each PLSR model is visualized in (A), while overlap for top negative VIPs is shown in (B). (C) Z-scored gene expression of the top 100 positive VIPs and top 100 negative VIPs for the cell count PLSR model. (D) Gene set enrichment analysis of the top VIPs for the Cell Count model. (E) Validation of Cell Count PLSR model using the DEPMAP Cancer Dependency Map (Project Achilles). Genes positively correlated with Cell Count show significant enrichment for essentiality in 94 breast cancer cell lines (χ² = 902.41, p-value < 2.2e-16). (F) Generalization of the Cell Count PLSR model to other cellular contexts. The model was applied to RNAseq data from 34 breast cancer cell lines and showed significant correlation (R² = 0.44, p-value < .001) with experimental proliferation rates.

We next focused on exploring and validating the Cell Count PLSR model, a well-studied phenotype that serves as an excellent test case of our approach to link image and molecular data. Genes positively correlated with cell count exhibited enrichment for gene sets canonically associated with proliferation, including mitotic cell cycle, DNA replication, and cyclin E2-CDK2 complex (**Figure 4C, D**). Among the positively correlated VIPs are essential components of mitosis, including several cell cycle checkpoint genes (CDK2, CDC6, CDCA4) (Chung & Bunz, 2010; Hayashi et al., 2006; Liu et al., 2000), microtubule regulation genes (TUBB, TUBB4B, TUBG1) (Giotti et al., 2019), and growth-regulating secreted factors (AREG, IL1A, CSF3) (Berasain & Avila, 2014; Chakraborty & Guha, 2007; Li et al., 2020). Conversely, the top negatively correlated VIPs associated with cell count were enriched for cellular component terms such as vacuolar lumen and lysosomal lumen, implying an upregulation of autophagy components in the absence of proliferation, potentially due to cellular stress (Mathiassen et al., 2017). These findings suggest that the Cell Count PLSR model captures expected biologically relevant pathways and processes associated with proliferation.

Assessing computational models using orthogonal approaches is crucial to ensure their robustness and reliability. Here, we leveraged independent, publicly available datasets to assess model generalizability and to validate our Cell Count model. First, we investigated whether the cell count signature identified genes essential for viability across a panel of diverse cancer cell lines. To achieve this, we utilized the Project Achilles dataset from the Cancer Dependency Map Portal (DEPMAP), which used high-throughput CRISPR Cas9 to experimentally determine gene essentiality across thousands of cancer cell lines (DepMap, 2024; Tsherniak et al., 2017). The Gene Effect score quantifies the impact of gene knockout on cell viability. The more negative the experimentally determined Gene Effect score, the greater its impact on viability. A Gene Effect score less than -0.5 indicates cell depletion, while a score less than -1 indicates strong cell killing. In contrast, a Gene Effect score of 0 signifies that a gene is not implicated in viability.

We examined the essentiality of the highest scoring VIPs included in our PLSR signature by analyzing their experimentally determined Gene Effect scores for the 94 breast cancer cell lines included in the DEPMAP database (**Figure 4E**). VIP genes that showed the highest positive correlation with cell count were significantly more likely to have a Gene Effect score ≤ -0.5 as compared to genes with negatively correlated VIP scores (χ² = 902.41, p-value < 2.2e-16). Moreover, across all genes included in our Cell Count PLSR model, those identified as high-scoring VIPs were more frequently associated with a Gene Effect score ≤ -0.5 than were genes with low-scoring VIPs (χ² = 3841.6, p-value < 2.2e-16), demonstrating the predictive capability of our model in assessing gene essentiality across diverse breast cancer cell lines (**Supp Fig 4E**). Notably, genes with high VIP scores in the PLSR model that also had large Gene Effect scores include canonical cell cycle components (CDC20, PLK1, GINS1) (Giotti et al., 2019) as well as recently discovered regulators (ALYREF) (Klec et al., 2022) and prognostic markers of breast carcinogenesis (GINS2) (Yu et al., 2020). This underscores that analyzing transcription and phenotype in a single cell line across multiple perturbations offers insights into mechanisms governing cell viability across diverse biological, while also revealing recently discovered genes involved in cell proliferation and potentially identifying novel, undiscovered ones. We also assessed the generalizability of the Cell Count PLSR model to predict proliferation in other cell line models. We analyzed publicly available datasets comprised of paired RNAseq and proliferation rates from 34 breast cancer cell lines (Heiser et al., 2012). For each cell line, we input the log2-normalized gene abundance data into our model to predict cell count. The model’s predicted cell count was significantly correlated with experimental measures of proliferation (R² = 0.44, p-value < .001) (**Figure 4F**). These results demonstrate that the Cell Count model generalizes across diverse cellular contexts beyond MCF10A and showcases the power of our approach to link molecular and image-based measurements. Taken together, these findings underscore the robustness of our approach in identifying functional molecular programs that govern complex phenotypic responses.

### Synergistic transcriptional upregulation of CXCR2 chemotactic signaling molecules via CREB activation promotes increased cell motility

To investigate the molecular mechanisms driving cell motility, we analyzed our Cell Motility model and the associated gene signature **(Figure 5A)**. Gene set enrichment analysis revealed that genes positively correlated with cell motility are significantly enriched in pathways related to responses to external and biotic stimuli, signaling receptor activator activity, and CXCR chemokine receptor binding (**Figure 5B**). To assess the generalizability of this model, we used the approach described above, but here leveraged our Cell Motility PLSR model to predict cell motility from publicly available RNAseq profiles from 28 breast cancer cell lines and compared this to experimentally determined migration rates (Heiser et al., 2012; Nair et al., 2019). The predicted cell motility values were strongly correlated with the experimentally measured migration rates, providing confidence that our model captures relevant biological programs driving cell motility (R² = 0.49, p-value < .001) (**Figure 5C)**.

**Figure 5:**
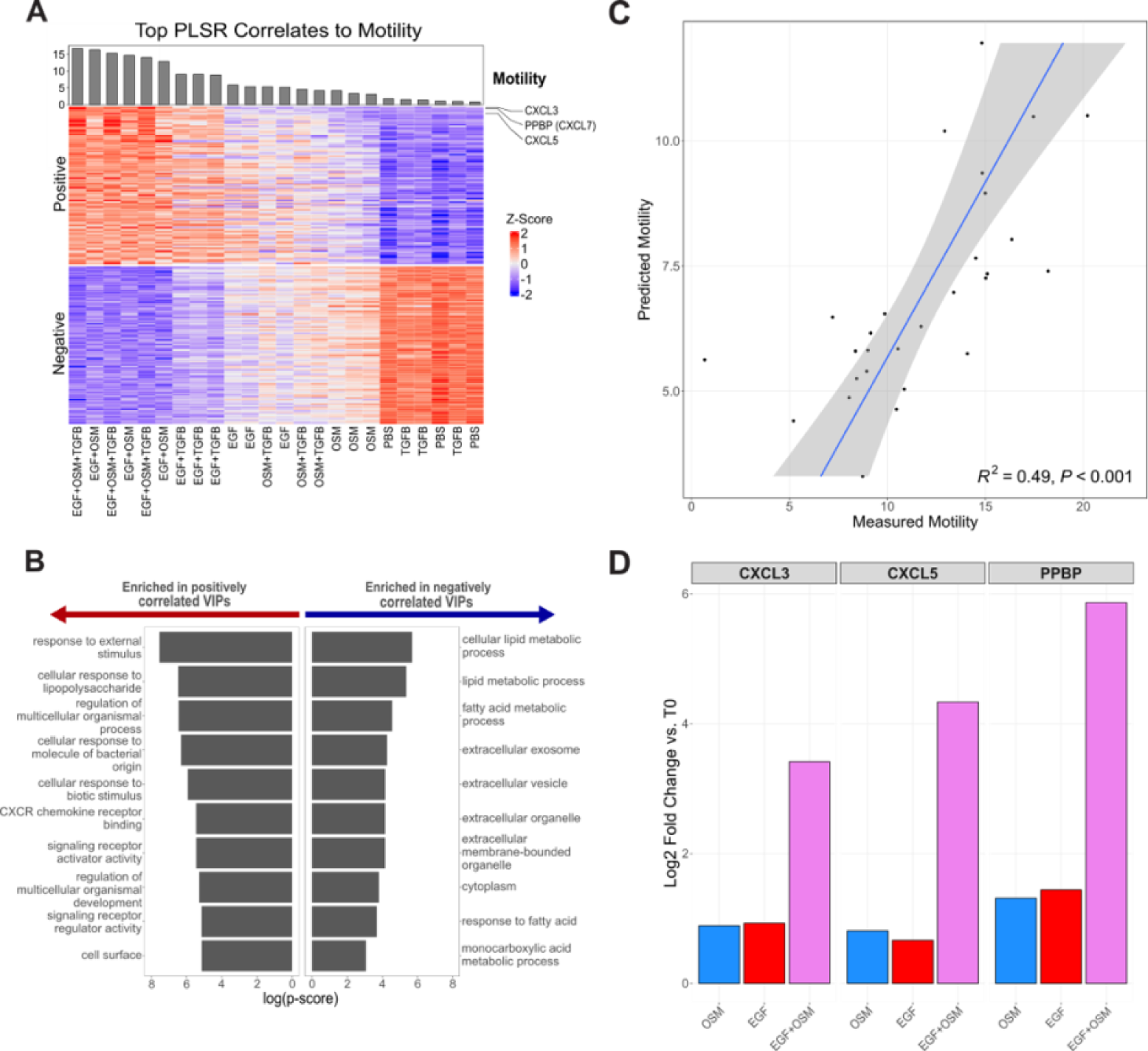
Expression of CXCR2 Agonists Correlates with Cell Motility. (A) Z-scored expression for the top 100 VIP genes positively and negatively with Cell Motility. (B) Gene set enrichment analysis of top VIPs, showing significant enrichment in pathways related to CXCR chemokine receptor binding. (C) Correlation between predicted cell motility values from the Cell Motility PLSR model and experimentally measured migration rates from 28 breast cancer cell lines. A significant correlation (R² = 0.49, p-value < .001) between predicted motility and experimental migration rates supports the relevance of the model in capturing motility-associated biological pathways. (D) LFC values compared to T0 control for the EGF+OSM combination condition and respective single ligand conditions show the synergistic upregulation of CXCR2 agonists (CXCL3, CXCL5, and PPBP).

Having established the validity of our Cell Motility model, we next more deeply explored it to uncover novel biological mechanisms driving this phenotypic response. Among the top most important genes are CXCL3, PPBP (CXCL7), and CXCL5, ranking first, second, and fifth in VIP scores (**Figure 5D**). These genes encode chemotactic ligands that signal through the CXCR2 chemokine receptor, a pathway known to enhance mammary epithelial cell migration (Singh et al., 2011). Furthermore, our previous RNAseq analysis showed that CXCL3, CXCL5, and PPBP are positively synergistically upregulated under the EGF+OSM condition, which produced the most significant increase in cell motility among all conditions tested **(Figure 1D**, **Figure 5D)**. Motivated by this, we focused on the EGF+OSM combination condition to functionally investigate the mechanism by which this combined treatment synergistically enhances motility as compared to the individual effects of EGF and OSM.

We hypothesized that the upregulation of CXCL3, CXCL5 and PPBP (CXCL7) contributes to the increased cell motility observed in the EGF+OSM condition compared to EGF and OSM single ligand conditions. To test this hypothesis, we first experimentally tested whether CXCR2 activation (the common receptor for CXCL3, CXCL5, and PPBP) influences cell motility in the EGF+OSM condition. We treated MCF10A cells with the ligand panel in the presence of AZD5069, a small molecule inhibitor of CXCR2 receptor activation, and then assessed cell motility. CXCR2 inhibition significantly suppressed cell motility in the EGF+OSM+TGFB, EGF+OSM, and OSM conditions, with the most substantial decrease observed in the EGF+OSM condition (23.9% median decrease across three biological replicates) **(Figure 6A, Supp Fig 5A)**.

**Figure 6:**
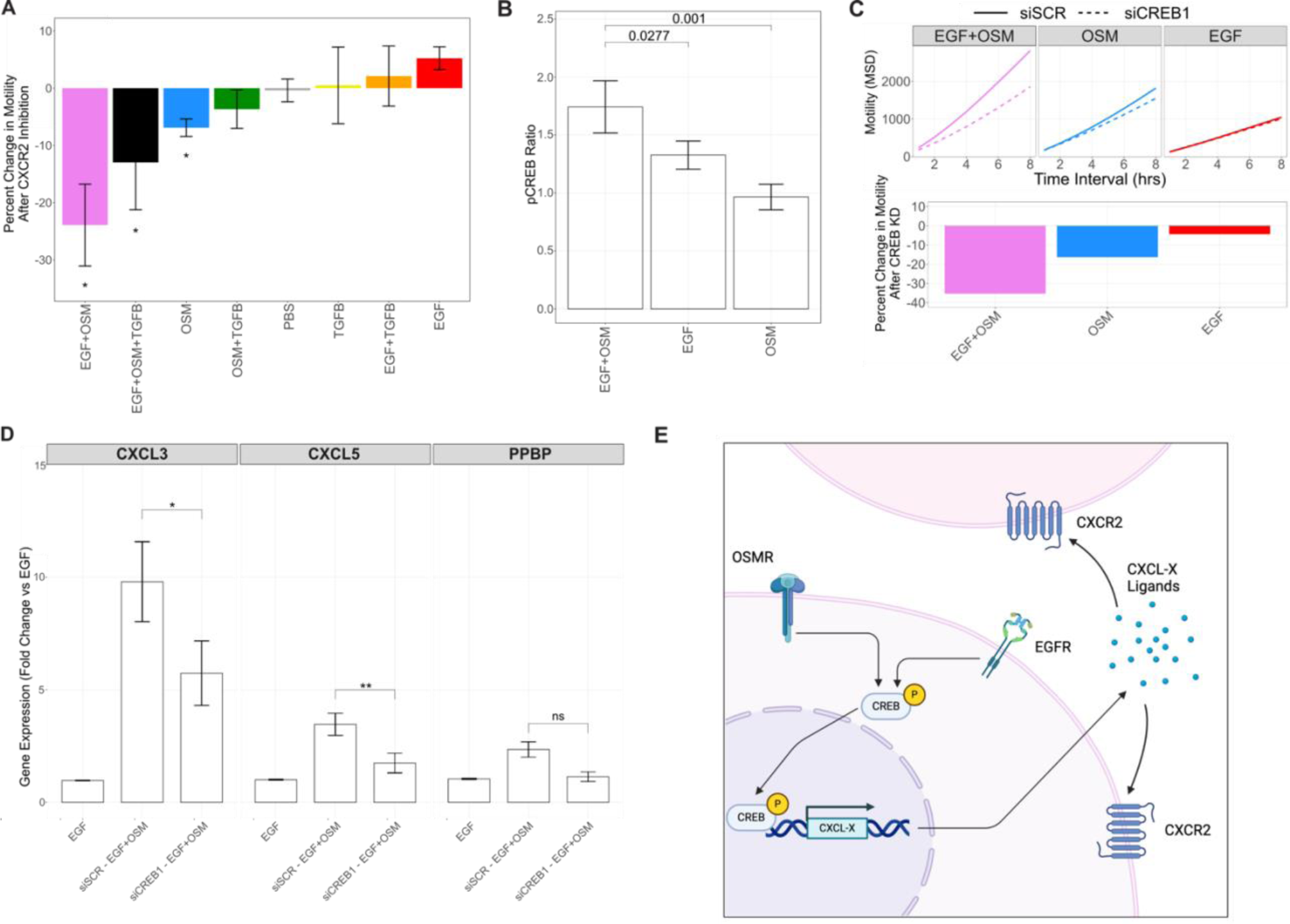
Synergistic Transcriptional Upregulation of CXCR2 Chemotactic Signaling Molecules via CREB Activation Promotes Increased Cell Motility. (A) Cell motility assays following treatment with single and combination ligand panel in the presence or absence of AZD5069, a CXCR2 inhibitor. CXCR2 inhibition significantly decreased cell motility in the EGF+OSM, EGF, and OSM conditions. Data shown as median change in motility with standard deviation from three biological replicates. P-value < .05 was considered significant. (B) Reverse Phase Protein Array (RPPA) analysis 1 hour after treatment with EGF, OSM, and EGF+OSM. Statistically significant changes in protein expression (p-value < .05) were assessed using Dunnett’s test. (C) Effects of CREB knockdown on cell motility of cells treated with EGF, OSM, or EGF+OSM. Mean Squared Displacement (MSD) is shown in the top panel and change in motility is shown in the bottom. (D) qPCR analysis of chemokine expression in CREB knockdown cells following EGF+OSM treatment. Barplots depict fold change in expression compared to the EGF control. Student’s t-test was used to assess significance (p-value < .05). (E) Putative mechanism of synergistic activation of CREB in response to combined EGF and OSM stimulation drives the upregulation of CXCL3, CXCL5, and PPBP, leading to increased cell motility via CXCR2 activation.

We then sought to identify the transcriptional regulators involved in the synergistic upregulation of CXCL3, CXCL5, and PPBP in the EGF+OSM condition. Given that this upregulation is synergistic in the EGF+OSM condition, and considering the emergent increase in cell motility observed with this combination treatment compared to either ligand alone, we hypothesized that the activation of transcriptional regulators would also exhibit a synergistic pattern in this context. To identify transcriptional regulators synergistically activated by EGF+OSM, we assessed proteomic changes with Reverse Phase Protein Array (RPPA) analysis 1 hour after ligand treatment (Tibes et al., 2006). Statistical analysis of the RPPA data identified 3 proteins with statistically significant change (p-value < .05) in expression between the EGF+OSM condition and both single ligand conditions: P70 S6 Kinase (pThr389), S6 (pSer240/244), and CREB (pS133) **(Figure 6B, Supp Fig 5B)**. CREB is a transcription factor known to enhance CXCL3, CXCL5, and PPBP expression (Sun et al., 2008), and its upregulation is consistent with our hypothesis that enhancement of chemotactic signaling contributes to the increased motility in the EGF+OSM condition.

We functionally validated CREB’s role in motility by performing siRNA knock-down in the presence of EGF, OSM, and EGF+OSM treatments followed by assessment of cell motility. CREB inhibition had minimal impact on EGF treated cells, a modest impact on OSM treated cells, and the greatest impact on EGF+OSM treated cells, mirroring the effects of CXCR2 inhibition (**Figure 6C**). Knockdown of CREB was confirmed through qPCR **(Supp Fig 5C).** We next sought to confirm the relationship between CREB activation and chemokine expression via qPCR analysis of CREB knockdown cells. Under EGF+OSM treatment, the expression of CXCL3 and CXCL5 was reduced, confirming that CREB indeed regulates the expression of these chemotactic ligands (**Figure 6D, Supp Fig 5D**).

In summary, our findings reveal that EGF+OSM promotes a synergistic increase in cell motility compared to the single ligand conditions. RNAseq analysis revealed that CXCL3, CXCL5, and PPBP are strongly correlated to cell motility and synergistically upregulated under the EGF+OSM condition. Inhibiting CXCR2 significantly reduced cell motility, and RPPA analysis indicated a synergistic increase in CREB phosphorylation, which enhances the expression of CXCL3, CXCL5, and PPBP. CREB knockdown experiments confirmed its crucial role, demonstrating that CREB activation drives the upregulation of these chemokines. These results collectively suggest that the synergistic activation of CREB, in response to combined EGF and OSM stimulation, drives the transcriptional upregulation of key chemokines, thereby promoting increased cell motility **(Figure 6E).**

## Discussion

Cells operate within a complex microenvironment in which signals from the local environment are continuously changing. These signals are integrated by cells, impacting transcription and signaling processes and thus significantly affecting cellular functions. While prior research has primarily investigated how individual signals combine to affect phenotypic and transcriptional outcomes, often in the context of drugs and a limited number of phenotypes like cell survival and proliferation, this study seeks to expand these concepts to the more intricate realm of ligand interactions. Here we explored the interactions between the ligands EGF, OSM, and TGFB and their effects on molecular and cellular responses. By analyzing the outcomes of both single-ligand and combined treatments, we reveal how these signals drive distinct transcriptional programs that impact cell motility, proliferation, and differentiation. Our findings show that combining ligand treatments can lead to unexpected phenotypic behaviors. For example, certain ligand combinations resulted in greater cell counts (EGF+OSM, OSM+TGFB), enhanced motility (EGF+OSM, EGF+TGFB, EGF+OSM+TGFB, OSM+TGFB), and altered both cell morphology and spatial arrangement (EGF+TGFB, OSM+TGFB, EGF+OSM+TGFB) as compared to their single-ligand treatments. These results demonstrate that the combined effects of ligand treatments can exceed the sum of their individual parts, emphasizing the challenge of predicting phenotypic responses from single-ligand data alone.

Distinct signaling pathways activated by EGF, OSM, and TGFB each play unique roles in cellular responses(Herbst, 2004; Massagué, 2012; Tanaka & Miyahima, n.d.). Our RNA sequencing data reveal that combination treatments mimic the gene expression profiles of one or both single-ligand conditions. An additive model of single-ligand gene expression showed a strong correlation with gene expression patterns observed in combination treatments, with the magnitude of transcriptional changes aligning with corresponding phenotypic effects. Notably, gene programs associated with epithelial differentiation are downregulated after nearly all treatment paradigms tested, with the notable exception of TGFB. This finding is intriguing given TGFB’s well-studied role in epithelial-to-mesenchymal transition (EMT) (Xu et al., 2009). This suggests that EGF might be necessary for TGFB to trigger the conventional EMT in MCF10A cells, highlighting the dependence of well-studied ligand-induced phenotypes (e.g. EMT induced by TGFB), on the presence of other ligands in the microenvironment.

Moreover, we identified unique transcription factors enriched across all combination conditions, including regulators of epithelial differentiation such as APP, BACH1, and CTNNB1. The shared enrichment of these transcription factors suggests that the combination of these ligands converges on similar signaling pathways to influence differentiation state. Building on prior research showing that small degrees of transcriptional synergy can influence phenotypic synergy (Diaz et al., 2020), we utilized an HSA-based modeling approach to define synergistic transcriptional programs. This approach uncovered subtle but significant synergistic gene sets specific to each ligand combination. While most studies have focused on phenotypic synergy, extending these frameworks into molecular synergy provides valuable insights into the underlying mechanisms driving combined ligand effects. Our studies support the adoption of existing frameworks designed for drug-induced changes in viability to gain insights into complex phenotypic responses. With the advent of spatially-resolved assays (Bressan et al., 2023), we envision that these frameworks could be applied to a broad array of data types and biological questions.

The use of Partial Least Squares Regression (PLSR) to connect transcriptional data with phenotypic outcomes is a significant strength of this study. Identifying gene signatures linked to cell count, motility, spatial organization and cytoplasmic size offers insights into the molecular drivers of these processes. Validation of the PLSR model with publicly available datasets and the Cancer Dependency Map underscores the robustness and generalizability of our PLSR models, with potential applications for identifying critical regulators of cell proliferation and survival (DepMap, 2024; Heiser et al., 2012; Nair et al., 2019; Tsherniak et al., 2017). Furthermore, this approach holds significant promise for uncovering novel regulators of cell phenotype by linking previously uncharacterized genes to specific cellular behaviors. However, alternative machine learning approaches such as random forests or neural networks could be employed in future studies to capture more complex non-linear relationships between transcriptional data and phenotypic outcomes (Ali & Aittokallio, 2019; Güvenç Paltun et al., 2021; Lind & Anderson, 2019; Torkamannia et al., 2023). These methods may offer complementary insights and help further refine our understanding of the complex signaling networks governing cellular behavior. Additionally, expanding the analysis to include time-course transcriptional data could provide a dynamic view of how these programs evolve over time in response to ligand treatments.

Our analysis of CXCR2 chemotactic signaling and CREB activation provides important mechanistic insights into molecular drivers of cell motility. The synergistic upregulation of CXCL3, CXCL5, and PPBP in the EGF+OSM condition, along with the effects of CXCR2 inhibition and CREB knockdown, highlights a key signaling axis involved in cell motility. CREB’s role in promoting chemokine expression further clarifies the molecular mechanisms underlying the observed changes. We demonstrated that EGF+OSM, only when applied in combination, phosphorylates and activates CREB transcription factor, leading to the transcriptional upregulation of CXCR2 receptor agonists. Activation of CXCR2 then contributes to increased cell motility.

Previous studies have established CREB’s role in promoting chemokine activity in malignant epithelial cells, transgenic mice, and other cell lines (Banerjee et al., 2011; Sun et al., 2008; Wen et al., 2013). While EGF is known to activate CREB in various contexts, including in breast tissue (Grinman et al., 2016), we reveal that in MCF10A cells, this pathway is uniquely activated by the combined presence of OSM and EGF. Additionally, prior research has shown that CXCR2 is overexpressed in breast cancer epithelial cells and enhances malignant cell migration (Bohrer & Schwertfeger, 2012; Miller et al., 1998). Here, we provide evidence for an autocrine signaling mechanism achieving similar outcomes in cell motility. However, it is important to note that the observed phenotypic changes likely also involve protein and signaling-driven mechanisms. While this study focused on the CREB-CXCR2 axis, future research should investigate other interacting pathways to provide a more comprehensive view of these regulatory networks on changes in motility. For instance, identifying which CXCR2 agonists actively bind to the receptor and exploring the roles of other transcription factors or co-regulators in chemokine expression and motility could offer deeper insights into these processes.

This study has several limitations that should be addressed in future research. First, while we focused on specific quantitative changes in cellular phenotype, many other aspects remain unexplored, such as metabolic activity, apoptosis, and differentiation status. Examining these phenotypes could provide critical insights into how ligand combinations influence cellular behavior. For instance, changes in metabolic activity could elucidate the energetic requirements for motility (Zanotelli et al., 2021), apoptosis assays might reveal how ligand signaling impacts cell survival (Gregory et al., 2016), and differentiation markers could help determine whether ligand combinations push cells toward specific lineages (Bussard & Smith, 2011). Additionally, investigating chromatin remodeling and transcriptional dynamics could uncover upstream regulatory mechanisms driving the observed phenotypic changes(Nava et al., 2019).

Second, our research was conducted using MCF10A cells, though we validated our results with external cancer cell datasets containing thousands of paired RNA-seq and phenotypic profiles to address this limitation. Future studies could expand the range of cell lines used, incorporating primary cells that more closely mimic the physiological state of cells in vivo (Richter et al., 2021). Additionally, using patient-derived organoids, a more complex model system that includes additional cell-cell contacts and extracellular matrix, could help determine the generalizability of our findings across different biological contexts (Rosenbluth et al., 2020).

Lastly, while our study observed various types of cell motility, we did not differentiate between distinct motility behaviors. Understanding the differences between random motility, directed migration, and collective movement could offer deeper insights into the regulation of these processes by ligand combinations (Trepat et al., 2012). To address this, future experiments could include chemotactic gradient assays to evaluate directed migration (Camley, 2018), scratch assays to study wound healing-like behavior (Martinotti & Ranzato, 2019), and 3D model systems to investigate the contributions of the ECM (Ravi et al., 2015). Such approaches would provide a more nuanced understanding of how specific signaling pathways influence distinct motility types.

Overall, this study offers a comprehensive analysis of how EGF, OSM, and TGFB signaling pathways interact to influence cellular responses through complex transcriptional programs. The integration of transcriptomic and phenotypic data using machine learning approaches enhances our understanding of the molecular mechanisms governing cell behavior, with potential implications for developing targeted therapeutic strategies to modulate cell motility and proliferation in cancer and other diseases.

## Methods

### MCF10A Cell Culture

Cell culture and ligand perturbation experiments were conducted as previously detailed (Gross et al., 2022). Briefly, for routine growth and passaging, cells were cultured in growth media containing DMEM/F12 (Invitrogen #11330-032), 5% horse serum (Sigma #H1138), 20 ng/ml EGF (R&D Systems #236-EG), 0.5 µg/ml hydrocortisone (Sigma #H-4001), 100 ng/ml cholera toxin (Sigma #C8052), 10 µg/ml insulin (Sigma #I9278), and 1% Pen/Strep (Invitrogen #15070-063). For perturbation experiments, growth factor-free media was used, composed of DMEM/F12, 5% horse serum, 0.5 µg/ml hydrocortisone, 100 ng/ml cholera toxin, and 1% Pen/Strep.

MCF10A cells were grown to 50-80% confluence in GM and detached using 0.05% trypsin-EDTA (Thermo Fisher Scientific #25300-054). Post-detachment, 5,000 cells were seeded into collagen-1 (Cultrex #3442-050-01) coated 24-well plates (Thermo Fisher Scientific #267062) in growth media. Six hours after seeding, cells were washed with PBS and growth factor-free media was added. After an 18-hour incubation in the new media, cells were treated with single ligand or combinations of ligands in fresh growth factor-free media: 10 ng/ml EGF (R&D Systems #236-EG), 10 ng/ml OSM (R&D Systems #8475-OM), and 10 ng/ml TGFβ (R&D Systems #240-B).

### Live-Cell Imaging and Image Analysis Pipeline

Live-cell imaging was performed using the Incucyte S3 microscope (Essen BioScience, #4647). Images were captured every 30 minutes for up to 48 hours. Live-cell image stacks were then registered using a custom Fiji script (Schindelin et al., 2012) and segmented with CellPose (Pachitariu & Stringer, 2022). Image tracking was carried out using the Baxter Algorithms pipeline (Magnusson et al., 2015).

All analysis of cell tracking data was performed in RStudio (R Core Team, 2022). The cell count metric was determined by counting the number of cells per field and normalizing this count by the T0 count for that field. Nearest neighbor distances were measured by calculating the pixel Euclidean distances from each cell centroid to the centroid of the second nearest cell in the imaging field. To account for variations in cell count, the mean nearest neighbor distances for each image were normalized by the expected mean distance to the nearest neighboring cell if the cells were distributed randomly (Ebdon, 1985). Cytoplasmic size was calculated as the average cytoplasmic size 24 hours after ligand addition. Cell motility was quantified by first removing tracks with distance jumps greater than 200 pixels in 30 minutes. Motility was estimated as the slope of the mean squared displacement (MSD) (Qian et al., 1991)over time intervals ranging from 30 minutes to 6 hours. The slope of the MSD for each treatment was derived by constructing a linear model comparing MSD to the time interval. This value is proportional to the diffusion coefficient for Brownian motion (Qian et al., 1991)

To assess the statistical significance of the deviation in combination ligand phenotypes from single ligand conditions, ANOVA was used followed by Tukey’s Honest Significant Differences test for post-hoc comparisons. For phenotype comparisons with EGF and PBS, ANOVA followed by Dunnett’s test for post-hoc comparisons was performed using the DescTools package in RStudio (Signorell & et mult. al., 2017). A p-value of less than 0.05 was considered significant for all tests.

### RNAseq Generation and Analysis

MCF10A cells were transferred into RLT Plus buffer (Qiagen) containing 1% β-ME, flash-frozen in liquid nitrogen, and stored at −80°C until RNA extraction. Total RNA was isolated from the frozen samples using the Qiagen RNeasy Mini Kit. cDNA libraries were prepared from poly(A)-selected RNA using the Illumina TruSeq Stranded mRNA Library Preparation kit. The Illumina HiSeq 2500 platform to generate 100-bp paired-end reads. Short read sequencing assays were performed by the OHSU Massively Parallel Sequencing Shared Resource.

RNAseq data was preprocessed and aligned using a pipeline adapted from Tatlow, et al (Tatlow & Piccolo, 2016). TrimGalore (v. 0.4.3) was used to trim adapter sequences and low-quality bases using CutAdapt (v. 1.10) and to generate FASTQ quality reports using FastQC (v. 0.11.5). After adapter trimming, reads were filtered to have a minimum length of 30 bp. Trimmed reads were pseudo-aligned to the GENCODE V24 transcriptome (GRCh38.p5) using Kallisto (v. 0.46.2). Gene-level quantifications were obtained from transcript-level abundance estimates using the R package tximport (v. 1.8.0) in R (v. 3.5.0).

We performed multiple differential gene expression analyses, comparing each ligand condition to time zero controls (**Figure 2A, B, D, E, F**) and comparing each two-ligand combination condition to the respective single ligand conditions (**Figure 3**). Both sets of analyses were conducted using RNA-seq gene-level summaries with the R package DESeq2 (version 1.24.0) (Love et al., 2014). For all analyses, significantly induced genes were defined as agenes with a LFC > 1.5 or LFC < -1.5 and a p-value < .05, adjusted using the Benjamini and Hochberg method (Benjamini & Hochberg, 1995).

Transcription factor enrichment scores were calculated using Priori with default settings, using TPM values as input (Yashar et al., 2024). Gene modules were identified through K-means clustering of the top 200 most up-regulated and top 200 most down-regulated genes, employing the ComplexHeatmap package (Gu et al., 2016). The number of clusters was determined by gap analysis. Statistical comparisons of differentially expressed genes within each module were performed using chi-squared analysis, followed by examination of standardized residuals. A p-value of less than 0.05 was considered significant. Gene set enrichment analyses were conducted using the Gprofiler package, focusing on Gene Ontology categories for Biological Process, Molecular Function, and Cellular Component (Aleksander et al., 2023; Ashburner et al., 2000; Kolberg et al., 2023). Gene sets were considered statistically significant if the adjusted p-value was below 0.01. The most enriched gene sets for each analysis were selected for visualization by ranking the gene sets first by the smallest p-value and subsequently by the highest odds ratio. P-value adjustments for multiple comparisons were made using the g:SCS method from the Gprofiler package ((Kolberg et al., 2023).

### Partial Least Squares Regression

PLSR models were built using replicate-level gene expression data as the independent variable to predict the replicate phenotypic metrics as the dependent variable. This pairing of replicates allowed us to utilize biological variation in both phenotypic and transcriptional responses. To exclude low-variance genes from the model, we used the VST method in the Seurat RStudio package (Version 3) to select the top 2,500 most variable genes for input (Stuart et al., 2019). Gene expression for each condition was normalized to T0 and scaled, and phenotypic metrics were scaled as well. The PLSR models were constructed using the PLS package in RStudio (Mevik & Wehrens, 2007). The number of components in each model was determined by identifying the elbows in the relative root mean squared error plots. Leave-one-out analysis was conducted to assess robustness. Genes were ranked by importance in the model by calculating the Variable Importance in Projection (VIP) scores (Wold et al., 2001). The top 100 VIP scoring genes, either positively or negatively correlated with the first component of the model, were used as input for gene set enrichment analyses. The enrichment was performed using the Gprofiler package, focusing on Gene Ontology categories for Biological Process, Molecular Function, and Cellular Component (Aleksander et al., 2023; Ashburner et al., 2000; Kolberg et al., 2023).

### Partial Least Squares Regression Model Validation

Orthogonal validation was conducted for the PLSR models predicting cell count and motility. This involved analyzing publicly available datasets containing RNAseq data paired with proliferation rates from 34 breast cancer cell lines (Heiser et al., 2012) and motility estimates from 28 breast cancer cell lines (Nair et al., 2019). We input the log2-normalized gene abundance data from these cell lines into our models to predict cell count or motility for each line. Pearson correlation was then calculated to compare the predicted phenotypes to the experimentally determined metrics.

Additionally, we validated the PLSR model predicting cell count using the Project Achilles dataset for breast cancer cell lines from the Cancer Dependency Map Portal (DEPMAP) (DepMap, 2024; Tsherniak et al., 2017). We compared VIP scores to Gene Effect scores calculated by DEPMAP from CRISPR screens and investigated the statistical significance of the relationship between VIP and Gene Effect scores using chi-squared analysis.

### CXCR2 Inhibition

Cells were cultured in 24-well plates following previously established protocols. After cell attachment and subsequent culture in assay media, 5 nM AZD5069 (MedChemExpress, #19855) or DMSO was added along with the ligand treatments. Cell imaging and motility assessments were performed as previously described. To determine the statistical significance of changes in motility, we first fitted an ordinary least squares linear model to the data using the estimatr package in RStudio (Blair et al., 2024). Then, we estimated marginal means with the emmeans package and computed pairwise contrasts to compare motility across all ligand conditions for inhibitor-treated versus DMSO-treated cells (Lenth, 2024). P-values were adjusted using Tukey’s method for multiple comparisons, and a p-value < 0.05 was considered significant.

### Reverse Phase Protein Array Sample Preparation

Cells were lysed and collected by manual scraping into 50-100 µL of lysis buffer (1% Triton X-100, 50 mM HEPES pH 7.4, 150 mM NaCl, 1.5 mM MgCl2, 1 mM EGTA, 100 mM Na pyrophosphate, 1 mM Na3VO4, 10% glycerol, 1x complete EDTA-free protease inhibitor cocktail (Roche #11873580001), 1x PhosSTOP phosphatase inhibitor cocktail (Roche #4906837001)). The lysates were incubated on ice for 20 minutes, followed by centrifugation at 14,000 rpm for 10 minutes at 4°C. The supernatant was collected, quantified using a BCA assay, and then mixed with 4X SDS sample buffer (40% glycerol, 8% SDS, 0.25 M Tris-HCl, 10% β-mercaptoethanol, pH 6.8). The mixture was boiled for 5 minutes and stored at -80°C. Three sets of biological replicates were submitted for RPPA testing. The samples underwent standard pre-processing using protocols established at the MD Anderson Cancer Center RPPA core (Akbani et al., 2014).

Statistical significance in antibody intensity between the EGF+OSM condition and both single ligand conditions was determined using ANOVA followed by Dunnett’s test using the DescTools package (Signorell & et mult. al., 2017). The RPPA data used in this study were part of a larger panel of conditions and perturbations assessed in MCF10A cells. The full dataset is available in **Supplementary Table 1**.

### SiRNA Assay and QPCR

Cells were seeded in growth media at a density of 25,000 cells per well in a 6-well plate. Seven hours after seeding, the cells were transfected with either a commercially validated siRNA pool targeting CREB1 (Horizon Discovery #L-003619-00-0005) or a negative control siRNA (Horizon Discovery #L-003619-00-0005) using Lipofectamine RNAiMAX Transfection Reagent (Invitrogen #13778100) at a concentration of 25 nM siRNA. After 48 hours, the cells were treated with EGF, OSM, or a combination of EGF and OSM for an additional 24 hours. Each treatment condition was performed in triplicate.

To evaluate the changes in mRNA levels following siRNA-mediated CREB knockdown, we extracted total RNA from treated cells using the RNeasy Mini Kit (Qiagen # 74104) as per the manufacturer’s protocol. We synthesized cDNA using the iScript cDNA Synthesis Kit (Bio-Rad #1708891). The mRNA expression levels were quantified through real-time qPCR with SYBR green chemistry on the Bio-Rad CFX Opus 384 Real-Time PCR System (Bio-Rad #12011452). The results were normalized to Glyceraldehyde 3-phosphate dehydrogenase (GAPDH) levels using the 2-ΔΔCt method (Livak & Schmittgen, 2001). Statistical significance between expression levels was determined by student’s T-test performed on ΔCt values (Student, 1908). The primer sequences for the target mRNAs are provided in **Supplementary Table 2**.

## Data Availability

Raw live-cell images of MCF10A cells treated with single and combination ligands are deposited on Zenodo.

RNA sequencing data and processed counts data generated for this study can be accessed from the Gene Expression Omnibus: GSE282654.

