## Supplemental Figures for "Integrative Analysis of EGF, OSM, and TGFB Signaling Pathways Reveals Synergistic Mechanisms Driving Cell Motility Through CXCR2 Chemotactic Signaling and CREB Activation"

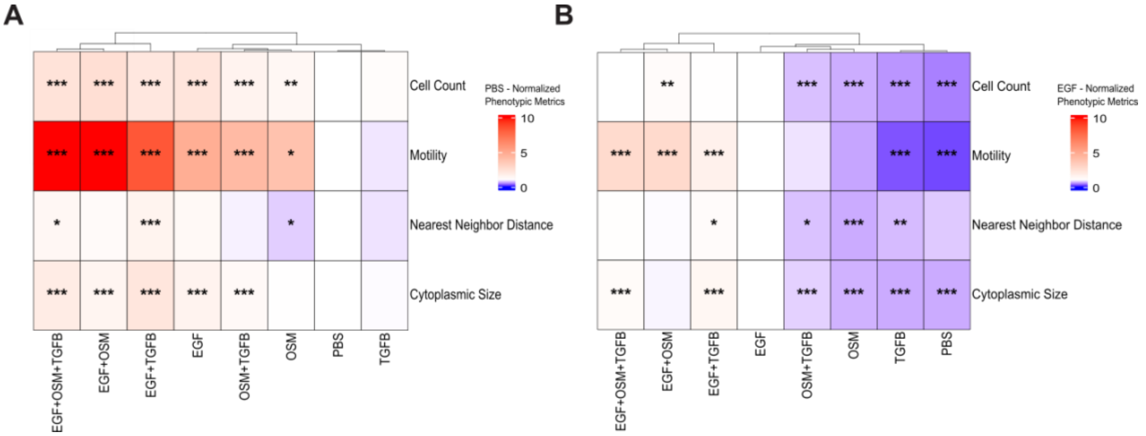


**Supp Figure 1: Statistical comparison of phenotypic responses to control conditions.**

(A) Quantified phenotypic metrics for all ligand treatments normalized by and compared to PBS. Statistical significance was determined using Dunnett's test with a p-value <.05 considered significant. (B) Quantified phenotypic metrics for all ligand treatments normalized by and compared to EGF. Statistical significance was determined using Dunnett's test with a p-value <.05 considered significant.

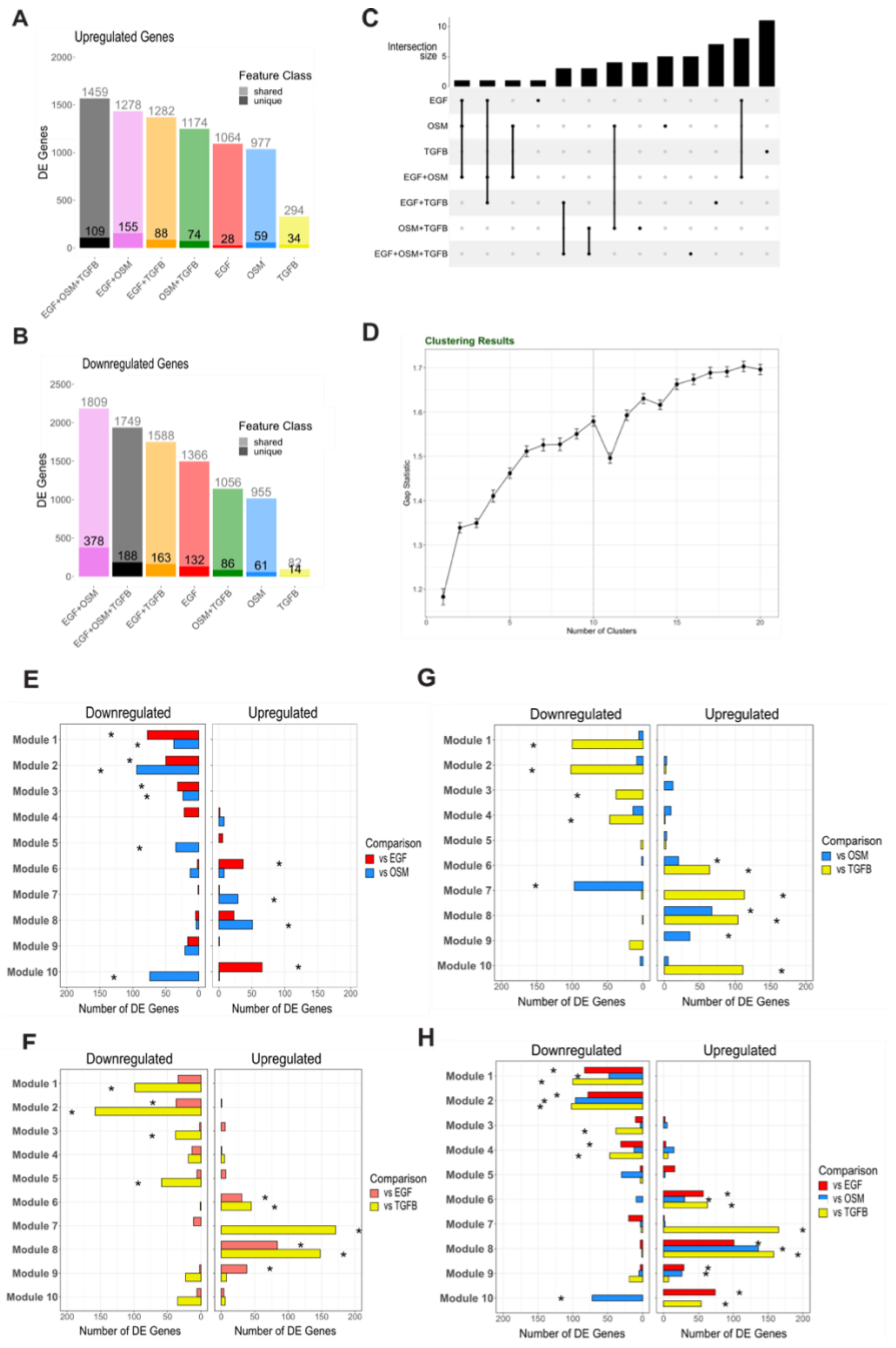


**Supp Figure 2: Detailed transcriptional analysis of single and combination ligand treatments.**

(A) Number of upregulated differentially expressed genes (LFC > 1.5, q-value < 0.05) for each treatment relative to PBS control. Genes unique to each treatment are shown with solid transparency, while those shared with at least one other condition are shown with lighter transparency. (B) Number of downregulated differentially expressed genes (LFC < -1.5, q-value < 0.05) for each treatment relative to PBS control, with transparency representing unique versus shared genes, as in panel A. (C) Upset plot showing the overlap of transcriptional regulators activated by single and combination treatments, highlighting the shared and unique regulatory programs across conditions. (D) Gap analysis identifies 10 optimal gene modules across treatments, based on clustering of transcriptional data. (E-H) Comparisons of gene module scores for each combination treatment versus the respective single ligand conditions. Statistical significance was determined by Chi-squared analysis, comparing the number of differentially expressed genes in each treatment relative to T0 for each module. P-value < .05 was considered significant.

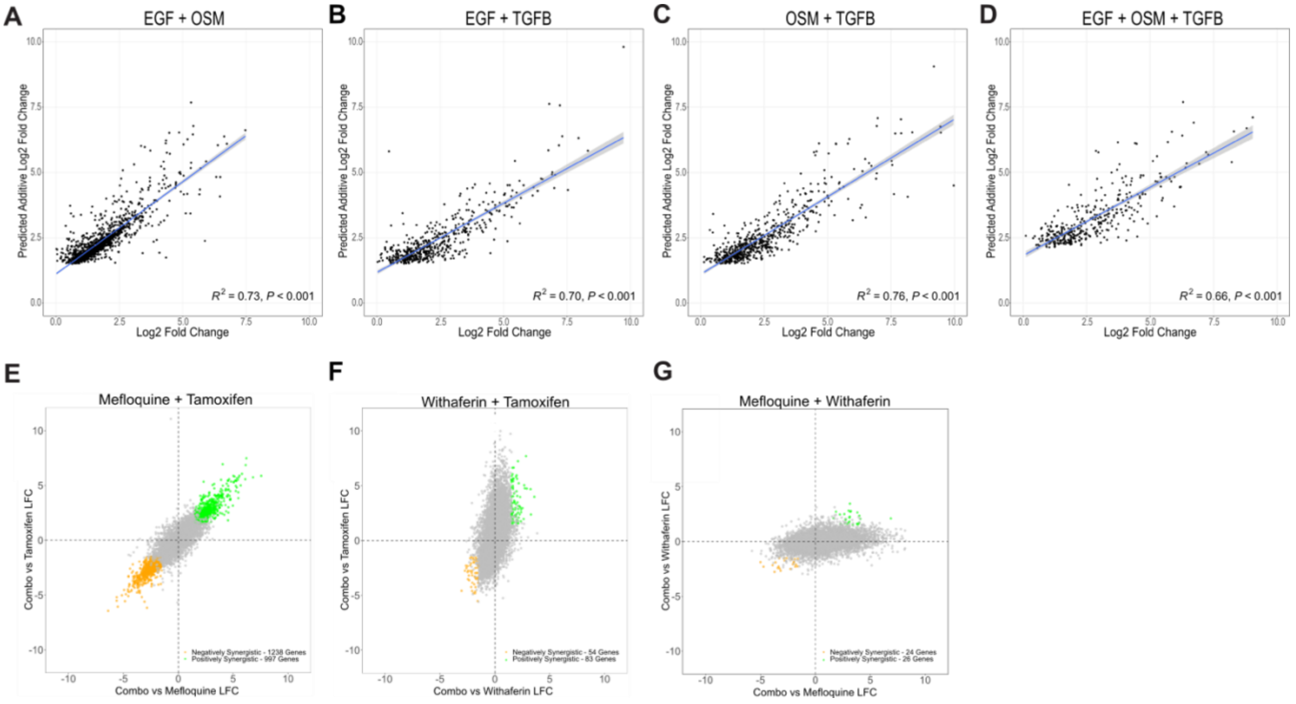


**Supp Figure 3: Additive modeling of ligand combinations and transcriptional synergy analysis of drug treatments.**

(A-D) Scatterplots depicting the correlation between the observed log2 fold change (LFC) for upregulated genes (LFC > 0.5) in each ligand combination condition (EGF+OSM, EGF+TGFB, OSM+TGFB, and EGF+OSM+TGFB) and the predicted LFC based on a purely additive model summing the LFC of the individual ligands. Pearson correlation coefficients ($R^2$) are reported for each comparison, demonstrating strong correlations ($R^2$ values ranging from 0.66 to 0.76, p-value < .001), indicating that ligand combination responses are predominantly additive. (E-G) Transcriptional synergy analysis applied to a published dataset of MCF7 cells treated with Tamoxifen, Mefloquine, and Withaferin individually and in combination. Scatterplots highlight the number of synergistic genes (positive and negative) induced by each drug combination, providing a comparison to ligand-induced transcriptional synergy.

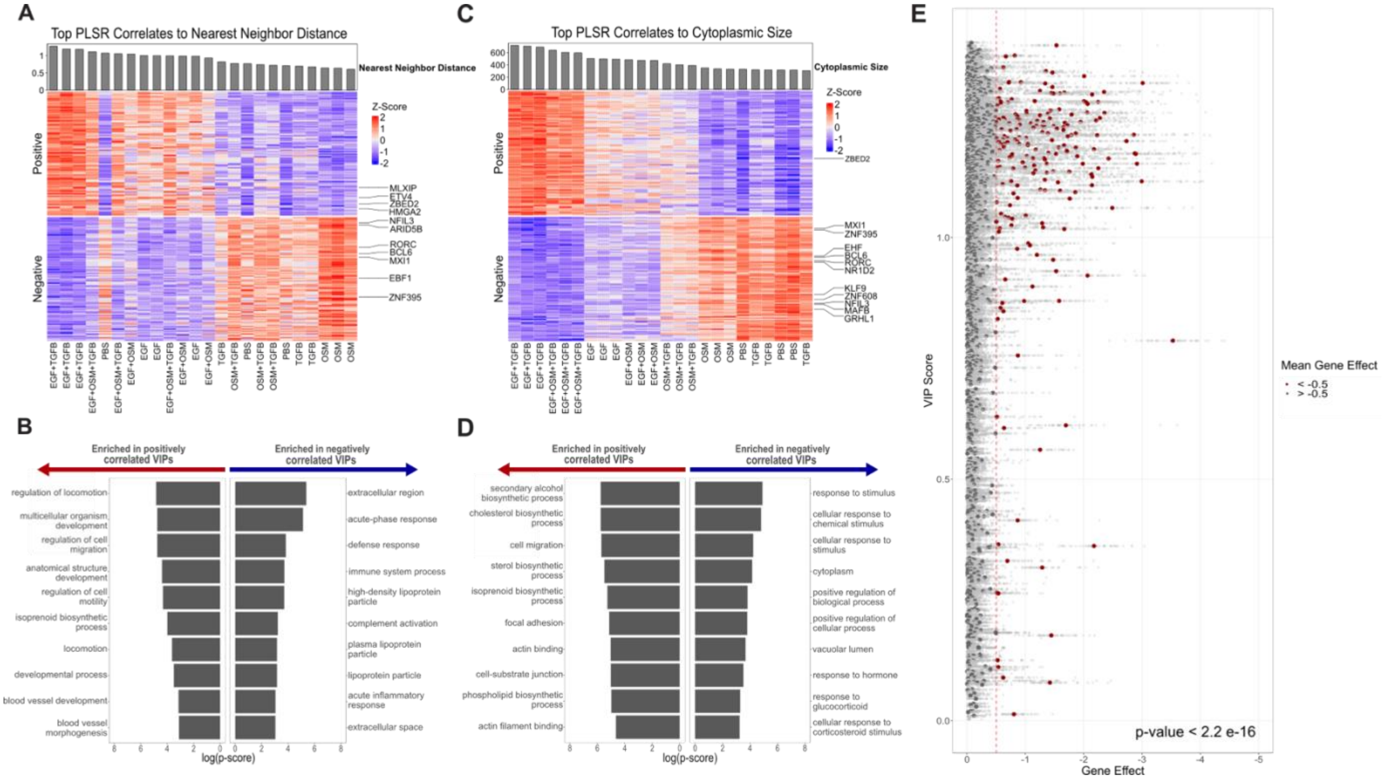


**Supp Figure 4: Gene Set Enrichment Analysis and Expression of VIP Genes from PLSR Models.**

(A) Z-scored expression of the top VIP genes associated with the Nearest Neighbor Distance phenotype in the PLSR model. (B) Gene set enrichment analysis of the top VIP genes associated with Nearest Neighbor Distance. (C) Z-scored expression of the top VIP genes associated with Cytoplasmic Size from the PLSR model. (D) GSEA of the top VIP genes associated with Cytoplasmic Size. (E) Evaluation of the Cell Count PLSR model using the DEPMAP dataset, where all genes in the model were analyzed for their Gene Effect scores across breast cancer cell lines. VIP genes that were positively correlated with cell count exhibited significantly lower Gene Effect scores ($\chi^2$ = 902.41, p-value < 2.2e-16), supporting their role in regulating cell viability and proliferation.

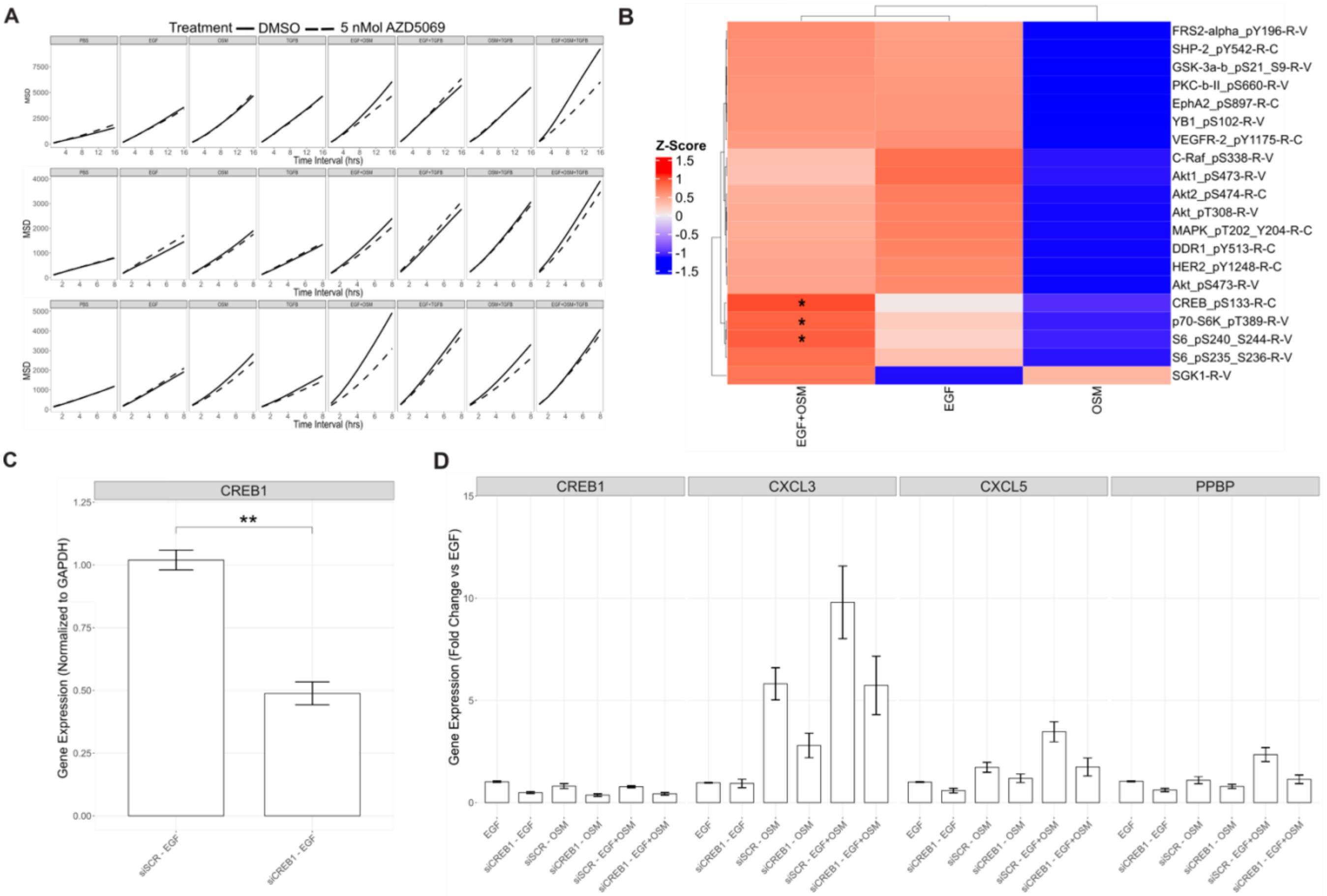


**Supp Figure 5: Cell Motility Assays, CREB Knockdown Efficiency, and RPPA Data.**

(A) Cell motility assay data showing the effects of CXCR2 inhibition (AZD5069) on MCF10A cell motility in the EGF+OSM, EGF, and OSM conditions, across all three biological replicates. The data depict the mean squared displacement, with CXCR2 inhibition significantly reducing motility in the EGF+OSM condition, (B) Reverse Phase Protein Array (RPPA) data from cells treated with EGF, OSM, and EGF+OSM. Dunnett's test was used to compare the single ligand conditions to the EGF+OSM combination, revealing statistically significant changes (p-value < .05) in CREB activation in the EGF+OSM condition. (C) qPCR confirmation of CREB knockdown in MCF10A cells, with treatment conditions for siRNA targeting CREB. (D) Full qPCR analysis of chemokine expression in CREB knockdown MCF10A cells treated with EGF, OSM, and EGF+OSM. The data show fold change of gene expression compared to EGF and normalized to GAPDH.
